# Illumina iSeq 100 and MiSeq exhibit similar performance in freshwater fish environmental DNA metabarcoding

**DOI:** 10.1101/2020.08.04.228080

**Authors:** Ryohei Nakao, Ryutei Inui, Yoshihisa Akamatsu, Masuji Goto, Hideyuki Doi, Shunsuke Matsuoka

## Abstract

Environmental DNA (eDNA) analysis is a method of detecting DNA from environmental samples and is used as a biomonitoring tool. In recent studies, Illumina MiSeq has been the most extensively used tool for eDNA metabarcoding. The Illumina iSeq 100 (hereafter, iSeq), one of the high-throughput sequencers (HTS), has a relatively simple workflow and is potentially more affordable than other HTS. However, its utility in eDNA metabarcoding has still not been investigated. In the present study, we applied fish eDNA metabarcoding to 40 water samples from river and lake ecosystems to assess the difference in species detectability and composition between iSeq and MiSeq. To check differences in sequence quality and errors, we also assessed differences in read changes between the two HTS. There were similar sequence qualities between iSeq and MiSeq. Significant difference was observed in the number of species between two HTS, but no difference was observed in species composition between the two HTS. Additionally, the species compositions in common with the conventional method were the same between the two HTS. According to the results, using the same amplicon library for sequencing, two HTS would exhibit a similar performance of fish species detection using eDNA metabarcoding.

## Introduction

Environmental DNA (eDNA) analysis methods can detect the DNA fragments shed from macro-organisms in environmental samples (water, sediment, or air). The eDNA methods can provide information on the distribution, abundance, seasonal change, and migration of species^1–4^ and can facilitate biodiversity monitoring activities^5,6^. Furthermore, eDNA methods use environmental samples for DNA detection, permitting non-invasive and non-destructive surveys in target species, habitats, and ecosystems^7^. One of the eDNA methods, eDNA metabarcoding, can detect multiple species from an environmental sample simultaneously using high-throughput sequencer (HTS)^8–11^. eDNA metabarcoding has been applied to detect both vertebrate^9,12–15^ and invertebrate^16–18^ compositions in communities. In addition, eDNA metabarcoding can detect higher levels of species diversity^19,20^ or complementary species diversity compared to conventional monitoring methods^10,21^.

Illumina MiSeq is the mainstream HTS for the detection species composition using eDNA metabarcoding^3, 9, 12, 16, 22^. In early 2019, the Illumina iSeq 100 (iSeq), which is a simpler and more affordable HTS system, was released^23^. The differences between iSeq and MiSeq are as follows, 1) base-calling systems, 2) the structure of the flow cell for loading the sequencing library, 3) the sequencing workflow^23^. The iSeq is simpler and requires less preparation using the cartridge, while the MiSeq requires relatively more preparation steps, including pre- and post-run wash of the flow channel. Such differences between the two sequencing approaches could influence species detectability and the sequencing quality during eDNA metabarcoding. Although some eDNA studies using iSeq have been reported ^24–26^, no comparative studies of sequencing performance and species detectability between iSeq and MiSeq have been performed in the eDNA metabarcoding.

In the present study, we applied fish eDNA metabarcoding using iSeq and MiSeq approaches to freshwater samples including river and lake ecosystems. We compared the sequence quality such as phred score and change rate of sequence read in pre-processing, and species detectability between iSeq and MiSeq. In addition, to evaluate the capacity of iSeq and MiSeq to detect species based on eDNA metabarcoding, we compared fish species compositions between eDNA metabarcoding (iSeq and MiSeq) and conventional methods.

## Results

### HTS using iSeq and MiSeq

In iSeq, the passing Filter (% PF), ≧ % Q30 (Read 1), and ≧ % Q30 (Read 2) were 80.80, 96.80, and 95.30%, respectively (Supplementary Fig. S1). In MiSeq, Passing Filter (% PF), ≧ % Q30 (Read 1), and ≧ % Q30 (Read 2) were 95.05, 97.30, and 96.48%, respectively (Supplementary Fig. S1). The % PF value of iSeq was slightly lower than that of MiSeq; however, this was due to differences in the %PF calculation methods^27^. The Q30 values of Read 1 and 2 were not remarkably different between iSeq and MiSeq.

In total, 3,325,177 and 2,154,367 read sequences were determined using iSeq and MiSeq, respectively. The sequencing depth and sequence per sample were consistent between iSeq and MiSeq in each processing step (Supplementary Table S2 and S3). From the results of spearman’s correlation tests, there were significant positive correlations between sequence reads of iSeq and MiSeq in merge pairs, quality filtering, and denoising step, respectively (ρ = 0.991, 0.991 and 0.993, *p* < 0.01, respectively; Table 1). Besides, there was significant positive correlation between the read ratio of iSeq and MiSeq after pre-processing (ρ = 0.993, *p* < 0.01; Table 1).

**Table 1.** Spearman’s correlations (ρ) on the number of sequence read per sample in each pre-processing step between iSeq and MiSeq

| Pre-processing step | Correlation ($\rho$) |
| --- | --- |
| Merge pairs | 0.991<sup>**</sup> |
| Quality filtering | 0.991<sup>**</sup> |
| Denoising | 0.993<sup>**</sup> |
| Taxonomic assignment | 0.993<sup>**</sup> |

<sup>\*\*</sup>: Correlation test, $p < 0.01$

### Taxonomic assignment of sequence read

Sequence reads obtained from iSeq and MiSeq in the river and lake samples after pre-processing are listed in Table 2. After the denoising step, most sequence reads in iSeq and MiSeq could be taxonomically assigned to fish species. In total, 154 and 168 representative iSeq and MiSeq sequences, respectively, were assigned to fish species (Assigned Reads in Table 2). After the taxonomic assignment, 102 and 101 freshwater fish sequences with ≧ 99 % identity were retained in iSeq and MiSeq, respectively. Low numbers of sequence reads of two species (*Rhinogobius* sp. and *Tridentiger* sp.) were retained from negative control samples (NC41–43) of iSeq and MiSeq (Supplementary Table S4 and S5), which could be due to cross-contamination among samples. The species were commonly observed throughout the samples, and the source of contamination could not be identified. Therefore, we did not assess cross-contamination across the samples using negative control samples.

**Table 2.** Ratio of sequence reads and operational taxonomic unit (OTU) count after denoising

|  |  | iSeq |  | MiSeq |  |
| --- | --- | --- | --- | --- | --- |
|  | Read Ratio |  |  |  |  |
|  | Denoised Read | 3010341 | 100.0% | 1914412 | 100.0% |
|  | Assigned Read | 2732599 | 90.8% | 1687629 | 88.2% |
|  | OTU Ratio |  |  |  |  |
|  | Total OTUs | 154 | 100.0% | 168 | 100.0% |
|  | OTU(Freshwater) | 102 | 66.2% | 101 | 60.1% |
|  | OTU (Marine and Brackish) | 24 | 15.6% | 22 | 13.1% |
|  | Under 99 % Identity | 28 | 18.2% | 45 | 26.8% |

Based on the sequencing results, retained iSeq and MiSeq sequences were assigned to 69 and 68 freshwater fish species, subspecies, or genera, respectively (Supplementary Table S4 and S5). The genera that were assigned to multiple candidate species are listed in Supplementary Table S6. To assess differences in species composition between iSeq and MiSeq, we used NMDS ordination, and the fish communities of iSeq and MiSeq at each site were plotted at almost similar coordinates (Fig. 1 (a)). From the results of PERMANOVA and PERMDISP, no differences of fish species compositions per samples were observed between iSeq and MiSeq (PERMANOVA, *p =* 0.95940 and PERMDISP, *p* = 0.51480 in Table 3). Sixty-eight fish species detected in MiSeq were also detected in iSeq and Japanese striped loach *Cobitis biwae* typeB was detected in L11 only from iSeq. The number of species per sample ranged from 4 to 27, in iSeq, and 4 to 26, in MiSeq. The differences in the number of species between iSeq and MiSeq ranged from 0 to 4. However, species that were detected by iSeq only often had low read counts (11 to 32 reads per species, Supplementary Table S7). For example, 15 and 12 species were detected in the iSeq and MiSeq at R5; however, the numbers of reads of the three species detected only in the iSeq were 11 (*Lepomis macrochirus*), 11 (*Oncorhynchus masou* subsp.), and 28 (*Tachysurus nudiceps*), respectively (Supplementary Table S7). All species detected only in iSeq remained after adjusting the number of sequence reads between iSeq and MiSeq using rarefaction at each site. For some species, the number of sequence reads were below the cut-off value (10 reads) for the pre-processing step after the rarefaction (Supplementary Table S7). For example, at R5, number of sequence read for *L. macrochirus* and *O. masou* subsp. were below the cut-off value, while that for *T. nudiceps* was above the cut-off value.

**Table 3.**
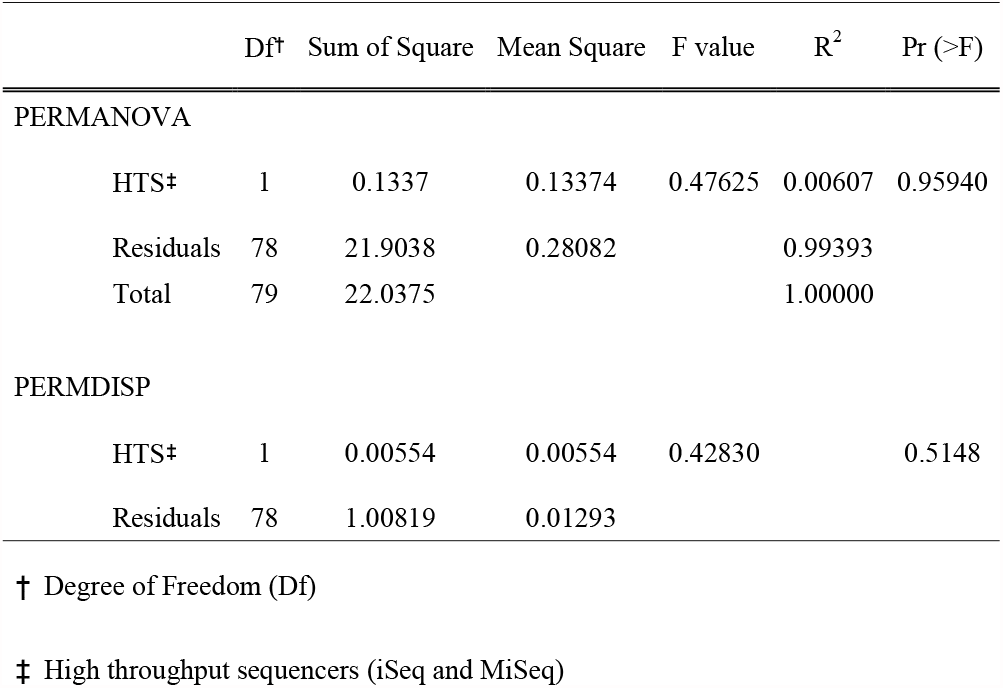
Statistical results of Permutational Multivariate Analysis of Variance (PERMANOVA) and Permutational Multivariate Analysis of Dispersion (PERMDISP) for comparisons of species composition between iSeq and MiSeq

**Figure 1.**
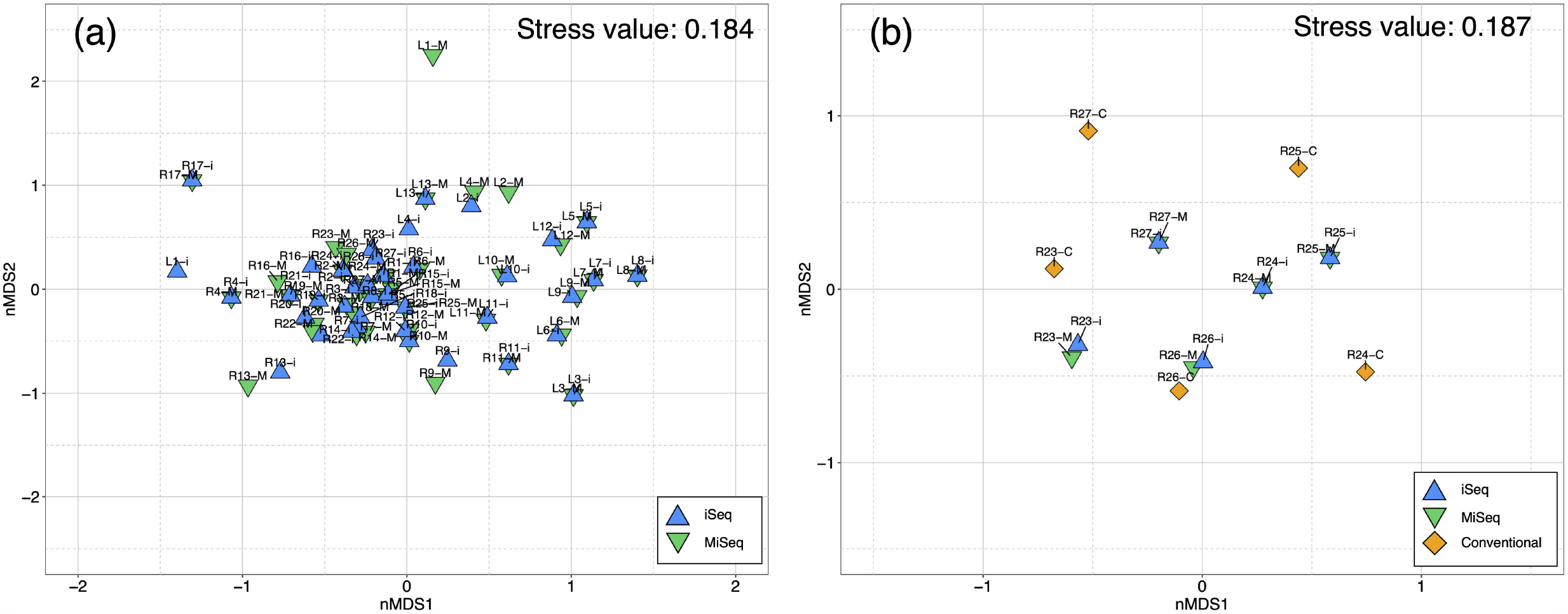
Two-dimensional NMDS plot of fish community using lake and river samples. (a) fish community of lake and river samples using iSeq and MiSeq. (b) fish community of five river samples using iSeq, MiSeq, and conventional methods. Blue triangles, green inverted triangles, and diamond shapes indicate the points of iSeq, MiSeq, and conventional methods, respectively. Labels with these shapes also indicate iSeq (-i), MiSeq (-M), and conventional methods (-C). The NMDS plots are performed based on incidence-based Jaccard index. Figure was created using “ggplot2” package version 3.3.0. supported on R program software^45^.

### Comparison of fish species composition between eDNA methods and conventional methods

R23–27 river samples were used for the comparisons between the two sequencers and the conventional methods (Fig. 2). A total of 30, 30, and 29 species were detected in R23–27 using the iSeq, MiSeq, and conventional methods, respectively (Supplementary Table S8). The number of species detected by iSeq, MiSeq, and conventional methods were 14–19, 14–19, and 8–16 in each site in R23–27, respectively (Supplementary Table S8). The number of species detected by iSeq and MiSeq was higher than that detected the conventional methods in all survey sites (deep green in Fig. 2). Based on the results of repeated measured ANOVA, there were significant differences among methods (F = 9.061, *p* = 0.0088), among sites (F = 4.827, *p* = 0.0282 in Supplementary Table S9). However, based on the results of the subsequent Tukey-Kramer test, there were no significant differences among the three methods (*p* > 0.05, Supplementary Table S10).

**Figure 2.**
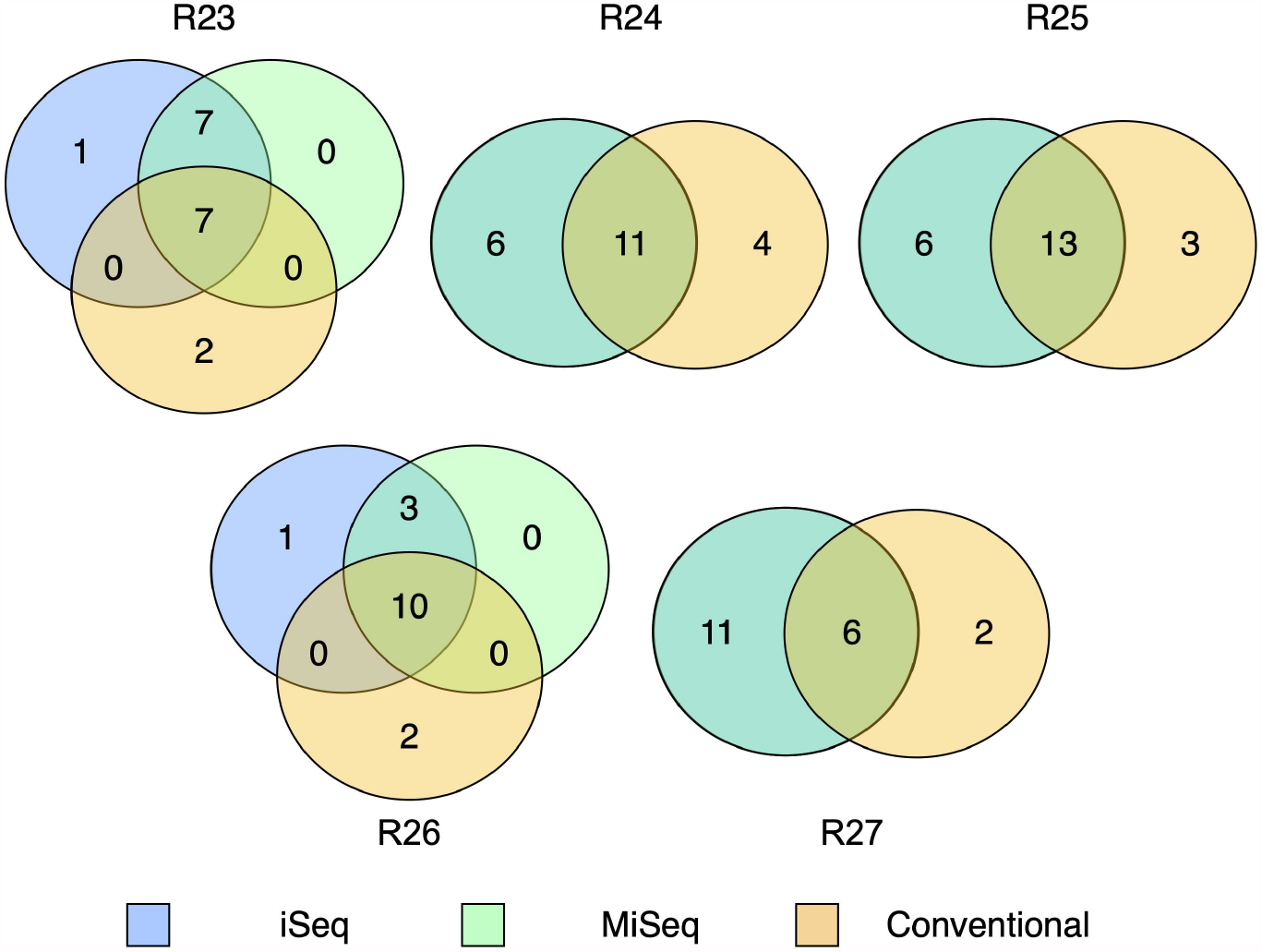
Venn diagrams for comparison of the number of species between high throughput sequencers and conventional methods in 5 river samples. Blue, green, and orange circles indicate the species of eDNA of iSeq, eDNA of MiSeq, and conventional methods, respectively. Deep green circles in R24, 25, and 27 indicate the species of two sequencers because of fully-overlapping species composition between iSeq and MiSeq. Note that the size of each circle does not represent a difference in the number of species. Figure was created using “venndiagram” package version 1.6.2 supported on R program software^44^.

Fish species compositions detected by HTS or captured by conventional methods in each river are shown in Figure 3. Some fish species were detected only in HTS through the five sites (e.g., *Pseudorasbora parva* and *Silurus asotus*). Three fish species, *Cobitis* sp. BIWAE type A, *Pseudobagrus aurantiacus* and *Oryzias latipes*, were observed only in the conventional methods (Fig. 3). However, other species observed in the conventional methods were also detected in HTS. No similarities were observed among fish species observed only in conventional methods at each site. To assess differences in species composition among the three methods, we used NMDS ordination (Fig. 1 (b)). In NMDS ordination, the fish communities of iSeq and MiSeq at each site were plotted at almost similar coordinates. Based on the results of PERMANOVA and PERMDISP analyses, there were no significant differences in species composition evaluated by iSeq, MiSeq, and conventional methods (PERMANOVA, *p =* 0.48450 and PERMDISP, *p* = 0.18510 in Table 4).

**Table 4.**
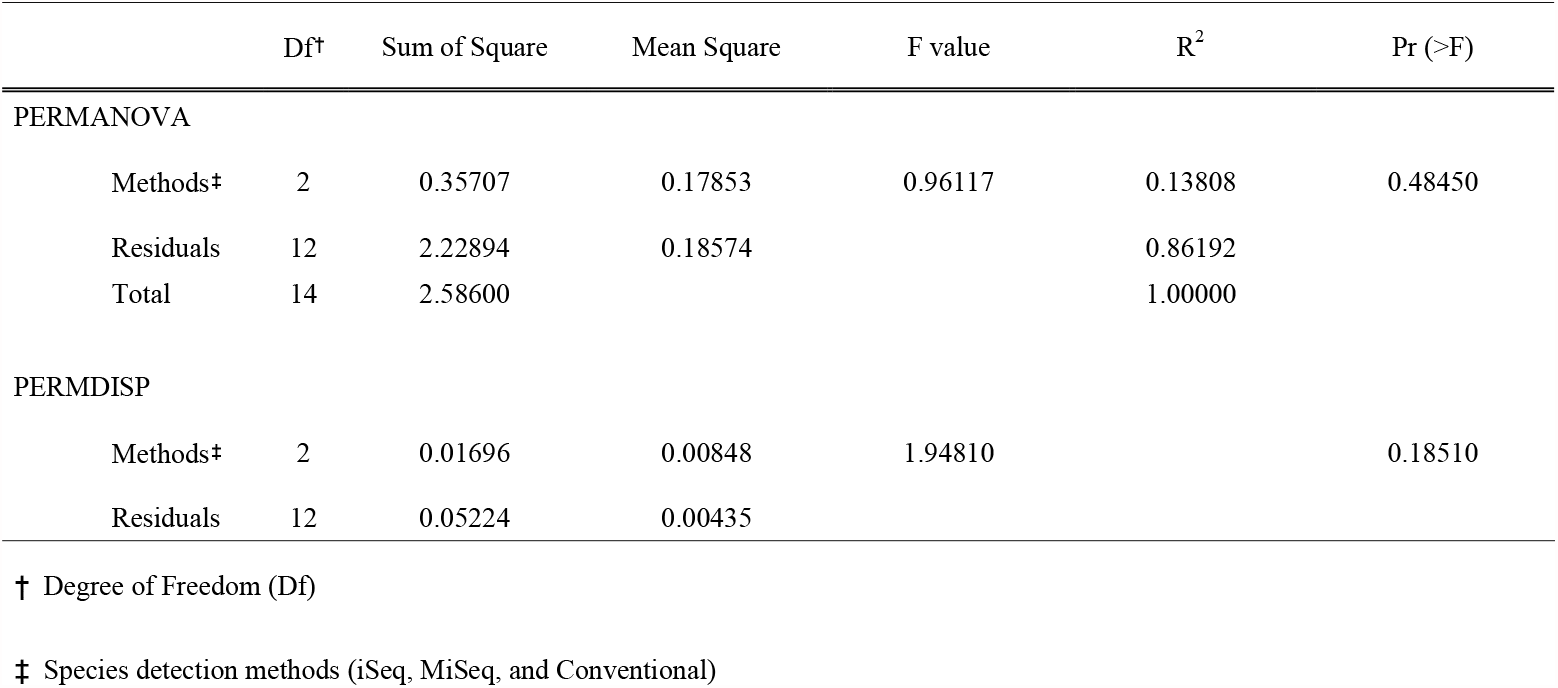
Statistical results of Permutational Multivariate Analysis of Variance (PERMANOVA) and Permutational Multivariate Analysis of Dispersion (PERMDISP) for comparisons of species composition among iSeq, MiSeq, and conventional methods

**Figure 3.**
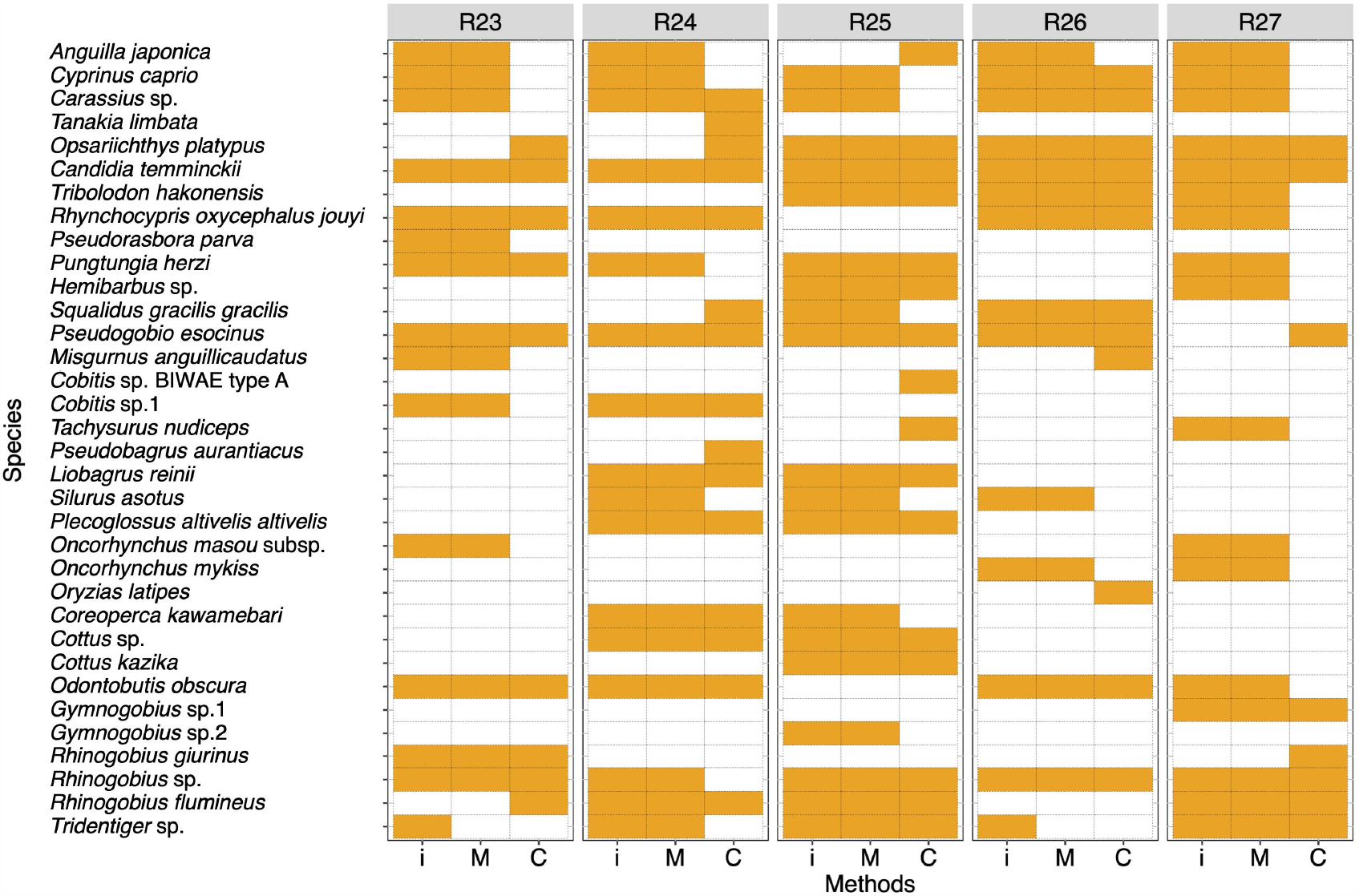
Heat map for comparison of the species compositions between high throughput sequencers and conventional methods in 5 river samples. Orange and white cells show the detection and non-detection of the species in each method. Abbreviations of “i”, “M”, and “C” indicate iSeq, MiSeq, and Conventional methods, respectively. Figure was created using “ggplot2” package version 3.3.0. supported on R program software^45^.

## Discussion

Here, we observed that iSeq and MiSeq could obtain similar sequence qualities and fish fauna in eDNA metabarcoding. As mentioned previously, the SBS chemistry of iSeq is distinct from that of MiSeq in sequencing workflow and base-calling, as well as distinct cartridge structures and flow cell mechanisms. However, such differences would not influence sequence quality in eDNA metabarcoding activities. From the results of two sequencing, high % PF and ≧ Q30 values in R1 and R2 have been obtained from the iSeq and MiSeq, respectively (Supplementary Fig. S1). The % PF value of iSeq was slightly lower than that of MiSeq (80.8% vs. 95.05%). In the iSeq, because template generation and the associated preliminary filtration steps are not applied, which leads to lower %PF values compared with MiSeq. Although % PF is much lower with the iSeq, it will not affect performance or data quality^27^. In addition, we obtained similar quality of sequence data from iSeq and MiSeq, and a few sequencing errors were observed in both HTS (Supplementary Fig. 1). Furthermore, the rate of change in the number of sequence reads in each process of the analytical pipeline was almost similar between iSeq and MiSeq (Supplementary Fig. S2-5). These changes in each step were positively correlated between the two HTS (Table 1). These results indicate that sequence qualities obtained from iSeq and MiSeq are similar and no sequence biases of the differences in sequencers are observed.

The freshwater fish fauna in the rivers and lakes identified by MiFish metabarcoding exhibited differences in the number of species or species composition between iSeq and MiSeq. A higher number of species per sample in iSeq than that in MiSeq could be due to the differences in the obtained sequence reads between the two sequencers. Species detected only in the iSeq had relatively low numbers of reads (11–32 reads, Supplementary Table S7). Sequence reads of these species remained after the rarefaction step, but some species have been fewer than 10 sequence reads after rarefaction (Supplementary Table S7). For example, at site 5, number of sequence read for *L. macrochirus* and *O. masou* subsp. were below the cut-off value, while that for *T. nudiceps* was above the cut-off value. At L4, all species detected only by iSeq after rarefaction (*Opsariichthys platypus, Candidia temminckii, Tribolodon brandtii*, and *Rhynchocypris oxycephalus jouyi*) were below the cut-off value. These species may have been eliminated by pre-processing before the taxonomic assignment. Therefore, for the same sequence depth, the difference in the number of species between iSeq and MiSeq will be much smaller. Such results indicate that the number of species that can be detected by iSeq and MiSeq are approximately similar if the same sequence library is used.

Based on the results of the comparisons of the numbers of species among the three methods, the number of species detected by eDNA metabarcoding was higher than those of conventional methods in each river, including many species that were only detected by HTS (Fig. 1 and Fig. 2). For many fish species detected only in HTS, hundreds to tens of thousands of sequence reads per species are obtained at each site and are unlikely to be potential contaminations from other samples (Supplementary Table S4 and S5). Therefore, these fishes are likely to be those that have been dropped by conventional methods because of the limitation of survey efforts (survey time and extent of survey area). Alternatively, the fish fauna of HTS may also contain the fish species inhabiting upstream from the sampling site and may reflect a wider range of fish fauna than conventional methods^29,30^. Some species were only observed in each river by conventional methods, but many of these species were also detected in HTS at other survey sites (e.g., *Anguilla japonica* in R25). Therefore, there was no common trend and ecological traits in fish species observed only by conventional methods. Of these species, three species (*Cobitis* sp. BIWAE type A, *Pseudobagrus aurantiacus*, and *Oryzias latipes*) have been observed only in conventional methods throughout five survey sites (Fig. 3). However, from the results of statistical analysis, there were no statistically significant differences in the numbers of observed species among the three methods. This statistical result could be due to the complementary effect of the number of species detected only by each method. The nMDS plot and statistical analyses also show that there was little difference in species composition among the three methods (Fig. 1 (b)). In previous studies, eDNA metabarcoding has exhibited “higher diversity” or “complementary” results when compared to the results of conventional methods^10, 11, 19^. Such complementary results in iSeq and MiSeq support the findings of previous studies.

Studies of eDNA metabarcoding have been increasing annually, and it is attracting the attention of researchers and stakeholders as a time- and cost-efficient method for detecting species composition and diversity^28^. Traditionally, MiSeq has been used extensively for eDNA metabarcoding^3, 9, 12, 16, 23^. Recently, some comparative studies using iSeq and other HTS, particularly MiSeq, have been reported in the medical field^31,32^. These studies show that iSeq can be used for whole-genome sequencing and targeted sequencing of pathogenic bacteria with accuracy similar to MiSeq. Some eDNA studies have been reported for the detection of fish community and aquatic plants, but have not compared the accuracy between iSeq and MiSeq^24–26^. Our results indicate that iSeq can be used for eDNA metabarcoding and has similar levels of species detectability to MiSeq. Despite the differences in sequencing depths, iSeq and MiSeq revealed similar fish fauna at each site. Furthermore, the differences in the number of species and species compositions between iSeq and MiSeq have been much smaller after the rarefaction. The results suggest that fish fauna from iSeq and MiSeq can be compared directly if library preparation is performed using similar processes. The iSeq has some limitations such as the unavailability of different kits with different lead lengths (max. 2 × 150 bp) and the number of leads (max. pair-end 4 million leads) compared to MiSeq. In contrast, the iSeq has fewer working procedures than MiSeq because of the use of cartridges and no need to clean the flow path. Therefore, iSeq may have lower cross-contamination risk between sequencing runs than MiSeq. Future research should evaluate cross-contamination risks such as the tag jumping rate among the Illumina indices and false-positive and false-negative detection of fish species between the two HTS. In addition, comparison of sequencing performance between iSeq and MiSeq should be also examined for eDNA libraries using other filtering methods (e.g., Sterivex and other materials).

## Materials and Methods

### Sample collection and filtration

We used 40 water samples for eDNA metabarcoding from 27 sites in 9 rivers and 13 lakes in Japan from 2016 to 2018 (Fig. 4). Sampling ID and detailed information for each site are listed in Supplementary Table S1. In the river water sampling, 1-L water samples were collected from the surface of at the shore of each river using bleached plastic bottles. In the field, a 1-ml Benzalkonium chloride solution (BAC, Osvan S, Nihon Pharmaceutical, Tokyo, Japan)^33^ was added to each water sample to suppress eDNA degeneration before filtering the water samples. We did not include field negative control samples in the HTS library, considering the aim of the presents study. The lake samples were provided by Doi et al. (2020)^34^ as DNA extracted samples. In the lake samples, 1-L water samples were collected from the surface at shore sites at each lake. The samples were then transported to the laboratory in a cooler at 4°C. Each of the 1-L water samples was filtered through GF/F glass fiber filter (normal pore size = 0.7 μm; diameter = 47 mm; GE Healthcare Japan Corporation, Tokyo, Japan) and divided into two parts (maximum 500-ml water per 1 GF/F filter). To prevent cross-contamination among the water samples, the filter funnels, and the measuring cups were bleached after filtration. All filtered samples were stored at -20□ in the freezer until the DNA extraction step.

**Figure 4.**
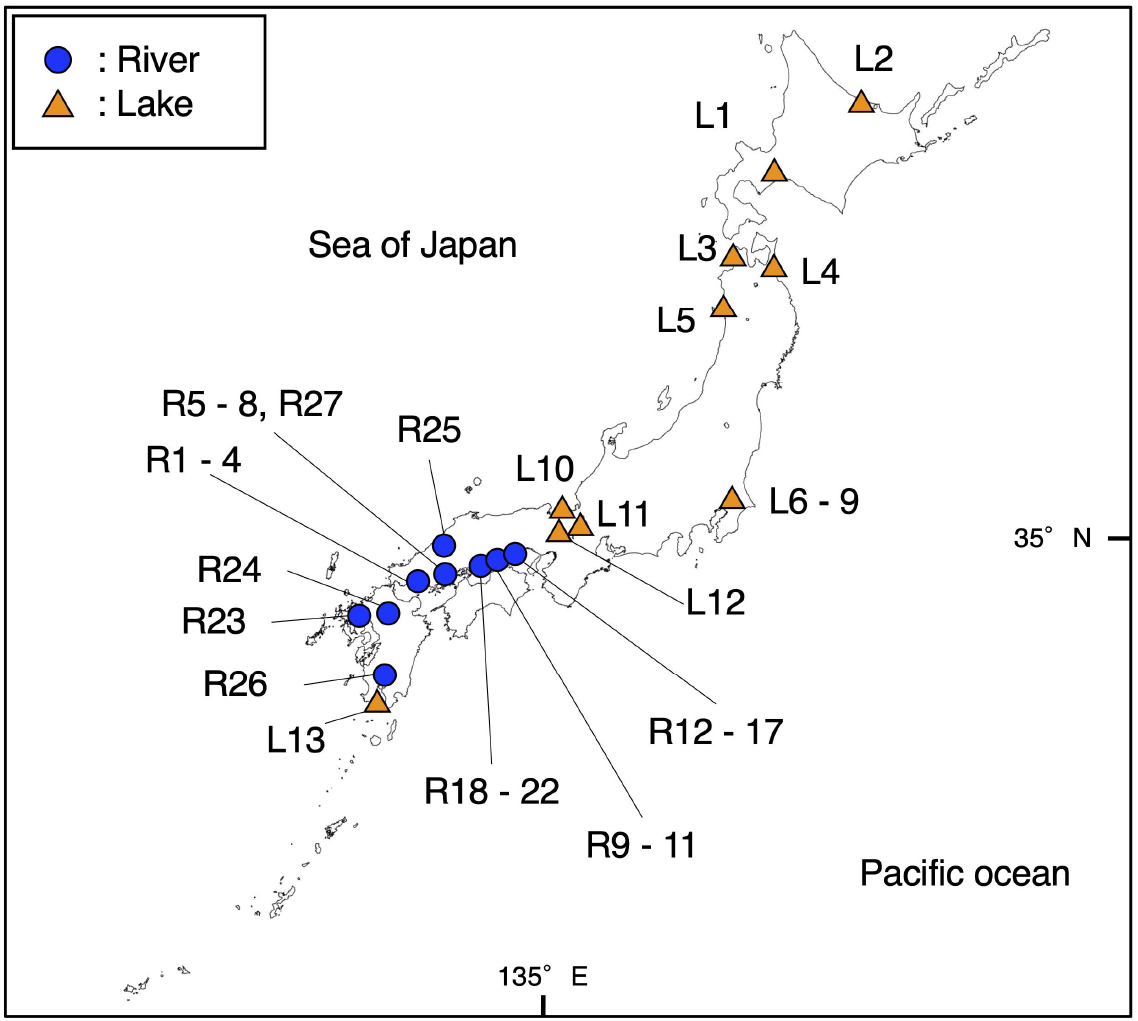
Sampling sites used in the present study. Blue circles and orange triangles show the locations of the river and lake samples, respectively. Detailed information on each site is listed in Supplementary Table S1. This map has been illustrated using QGIS ver.3.10 (http://www.qgis.org/en/site/) based on the Administrative Zones Data (http://nlftp.mlit.go.jp/ksj/gml/datalist/KsjTmplt-N03-v2_3.html) which were obtained from free download service of the National Land Numerical Information (http://nlftp.mlit.go.jp/ksj/index.html, edited by RN). There was no need of obtaining permissions for editing and publishing of map data.

### DNA extraction and library preparation

The total eDNA was extracted from each filtered sample using the DNeasy Blood and Tissue Kit (QIAGEN, Hilden, Germany). Extraction methods were according to Uchii *et al*.^35^, with a few modifications. A filtered sample was placed in the upper part of a Salivette tube and 440 μL of a solution containing 400 μL Buffer AL and 40 μL Proteinase K added. The tube with the filtered sample was incubated at 56°C for 30 min. Afterward, the tube was centrifuged at 5000 ×*g* for 3 min, and the solution at the bottom part of the tube was collected. To increase eDNA yield, 220-μL Tris-EDTA (TE) buffer was added to the filtered sample and the sample re-centrifuged at 5000 ×*g* for 1 min. Subsequently, 400 μL of ethanol was added to the collected solution, and the mixture was transferred to a spin column. Afterward, the total eDNA was eluted in 100-μL buffer AE according to the manufacturer’s instructions. All eDNA samples were stored at -20°C until the library preparation step.

In the present study, we used a universal primer set “MiFish” for eDNA metabarcoding^9^. The amplicon library was prepared according to the following protocols. In the first PCR, the total reaction volume was 12 μL, containing 6.0μL 2× KOD buffer, 2.4 μL dNTPs, 0.2 μL KOD FX Neo (TOYOBO, Osaka, Japan), 0.35 μL MiFish-U-F (5’-*ACACTCTTTCCCTACACGACGCTCTTCCGATCTNNNNNN*GTCGGTAAAACTCGTGCCA GC -3’), MiFish-U-R (5’-*GTGACTGGAGTTCAGACGTGTGCTCTTCCGATCTNNNNNN*CATAGTGGGGTATCTAAT CCCAGTTTG -3’), MiFish-E-F (5’-*ACACTCTTTCCCTACACGACGCTCTTCCGATCTNNNNNN*RGTTGGTAAATCTCGTGCC AGC -3’) and MiFish-E-R (5’-*GTGACTGGAGTTCAGACGTGTGCTCTTCCGATCTNNNNNN*GCATAGTGGGGTATCTAA TCCTAGTTTG -3’) primers with Illumina sequencing primer region and 6-mer Ns, and 2 μL template DNA. The thermocycling conditions were 94□ for 2 min, 35 cycles of 98□ for 10 s, 65□ for 30 s, 68□ for 30 s, and 68□ for 5 min. The first PCR was repeated four times for each sample, and the replicated samples were pooled as a single first PCR product for use in the subsequent step. The pooled first PCR products were purified using the Solid Phase Reversible Immobilization select Kit (AMPure XP; BECKMAN COULTER Life Sciences, Indianapolis, IN, USA) according to the manufacturer’s instructions. The DNA concentrations of purified first PCR products were measured using a Qubit dsDNA HS assay kit and a Qubit 3.0 fluorometer (Thermo Fisher Scientific, Waltham, MA, USA). All purified first PCR products were diluted to 0.1 ng/μL with H_2_O, and the diluted samples were used as templates for the second PCR. In the first PCR step, the PCR negative controls (four replicates) were included in each experiment. A total of three PCR negative controls were included in the library (PCR Blank 1–3 samples in Supplementary Table S1, S2, S4, and S5).

The second PCR was performed to add HTS adapter sequences with 8-bp dual indices. The total reaction volume was 12 μL, containing 6.0 μL 2× KAPA HiFi HotStart ReadyMix, 1.4 μL forward and reverse primer (2.5 μM), 1 μL purified first PCR product, and 2.2 μL H_2_O. The thermocycling conditions were 95□ for 3 min, 12 cycles of 98□ for 20 s, 72□ for 15 s, and 72□ for 5 min.

Each Indexed second PCR product was pooled in the equivalent volume, and 25 μL of the pooled libraries were loaded on a 2% E-Gel SizeSelect agarose gels (Thermo Fisher Scientific), and a target library size (ca. 370 bp) was collected. The quality of the amplicon library was checked using an Agilent 2100 Bioanalyzer and Agilent 2100 Expert (Agilent Technologies Inc., Santa Clara, CA, USA), and the DNA concentrations of the amplicon library were measured using Qubit dsDNA HS assay Kit using a Qubit 3.0 fluorometer.

### High-throughput sequencing

Amplicon library was sequenced using iSeq and MiSeq platforms (Illumina, San Diego, CA, USA). To normalize the percentage of pass-filtered read numbers, the sequencing runs using the same libraries were performed using iSeq i1 Reagent and MiSeq Reagent Kit v2 Micro. Both sequencing was performed with 8 million pair-end reads and 2×150 bp read lengths. Each library was spiked with approximately 20% PhiX control (PhiX Control Kit v3, Illumina, San Diego, CA, USA) before sequencing runs according to the recommendation of Illumina. The wells of cartridges in the iSeq run were loaded with 20 μL of 50 pM library pool, and sequencing performed at Yamaguchi University, Yamaguchi, Japan. The wells of cartridges for MiSeq runs were loaded with 600 μL of 16 pM library pool, and sequencing performed at Illumina laboratories (Minato-ku, Tokyo, Japan). Subsequently, the sequencing dataset outputs from iSeq and MiSeq were subjected to pre-processing and taxonomic assignments. All sequence data are registered in the DNA Data Bank of Japan (DDBJ) Sequence Read Archive (DRA, Accession number: DRA10593).

### Pre-processing and taxonomic assignments

We used the USEARCH v11.0667^36^ for all data pre-processing activities and taxonomic assignment of the HTS datasets obtained from the iSeq and MiSeq platforms^16, 37^. First, pair-end reads (R1 and R2 reads) generated from iSeq and MiSeq platforms were assembled using the “fastq_mergepairs” command with a minimum overlap of 10 bp. In the process, the low-quality tail reads with a cut-off threshold at a Phred score of 2, and the paired reads with too many mismatches (> 5 positions) in the aligned regions were discarded^38^. Secondly, the primer sequences were removed from the merged reads using the “fastx_truncate” command. Afterward, read quality filtering was performed using the “fastq_filter” command with thresholds of max expected error > 1.0 and > 50 bp read length. The pre-processed reads were dereplicated using the “fastx_uniques” command, and the chimeric reads and less than 10 reads were removed from all samples as the potential sequence errors. Finally, an error-correction of amplicon reads, which checks and discards the PCR errors and chimeric reads, was performed using the “unoise3” command in the unoise3 algorithm^39^. Before the taxonomic assignment, the processed reads from the above steps were subjected to sequence similarity search using the “usearch_global” command against reference databases of fish species that had been established previously (MiFish local database v34). The sequence similarity and cut off E-value were 99 % and 10^−5^, respectively. If there was only one species with ≧ 99 % similarity, the sequence was assigned to the top-hit species. Conversely, sequences assigned to two or more species in the ≧ 99 % similarity were merged as species complex and listed in the synonym group. Generally, the species complexes were assigned to the genus level (e.g., Asian crucian carp *Carassius* spp.). Species that were unlikely to inhabit Japan were excluded from the candidate list of species complexes. For example, the sequence of one of bitterling *Acheilignathus macropterus* included other different two species, *A. barbatus* and *A. chankaensis*, as the species of the 2nd hit candidate; however, the two species are not currently found in Japan. Therefore, the sequence was assigned to *A. macropterus* in the present study. Because we used only freshwater fish species, we removed the operational taxonomic units (OTUs) assigned to marine and brackish fishes from each sample. Finally, sequence reads of each fish species were arranged into the matrix, with the rows and columns representing the number of sites and fish species (or genus), respectively.

We evaluated sequence quality based on 1) the percentage of clustering passing filter (% PF) and 2) sequencing quality score ≧ % Q30 (Read1 and Read2) between iSeq and MiSeq platforms. The % PF value is an indicator of signal purity for each cluster^40^. The condition leads to poor template generation, which decreases the % PF value^40^. In the present study, a >80 % PF value was set as the threshold of sequence quality in iSeq and MiSeq runs. Sequence quality scores (Q score) measure the probability that a base is called incorrectly. Higher Q scores indicate lower probability of sequencing error, and lower Q scores indicate probability of false-positive variant calls resulting in inaccurate conclusions^41^. In the present study, the % Q30 values (error rate = 0.001 %) were used for the comparison of sequence quality between iSeq and MiSeq. The parameters were collected directly using Illumina BaseSpace Sequence Hub. We also evaluated changes in sequence reads in pre-processing steps between iSeq and MiSeq platforms. Sequence reads were assessed based 1) merge pairs, 2) quality filtering, and 3) denoising. In each step, the change in the number of reads before and after processing was calculated. The calculated numbers of sequence reads are listed in Supplementary Table S2 and S3 in series.

### Comparing sequence quality and fish fauna between iSeq and MiSeq

To test a relationship of remained sequence reads between iSeq and MiSeq in each pre-processing part, we performed spearman’s rank correlation test in each step. In the present study, however, the sequencing run by iSeq and MiSeq was performed only once each for the same sample. Therefore, we could not assess the variabilities of the sequence read in quality checks and taxonomic assignment in the same samples between iSeq and MiSeq.

Before the comparison of fish fauna, rarefaction curves were illustrated for each sample in both iSeq and MiSeq to confirm that the sequencing depth adequately covered the species composition using the “rarecurve” function of the “vegan” package ver. 2.5-6 (https://github.com/vegandevs/vegan) in R ver. 3.6.2^42^. In the present study, the differences in the numbers of sequence reads among samples were confirmed in the two sequencers, but rarefaction curves were saturated in all iSeq and MiSeq samples (Supplementary Fig. S6 and S7). We performed a rarefaction using the “rrarefy” function in “vegan” package to match up the iSeq sequence depths of each sample with that of MiSeq. However, the number of species in each sample on the iSeq have not changed before or after the rarefaction. Therefore, we have used the raw data set before the rarefaction for the subsequent analyses.

We compared the species detection capacities of iSeq and MiSeq based on environmental DNA metabarcoding. Using fish faunal data obtained from iSeq and MiSeq, non-metric multidimensional scaling (NMDS) was performed in 1000 separate runs using the “metaMDS” functions in the “vegan” package ver. 2.5-6. For NMDS, the dissimilarity of the fish fauna was calculated based on the incidence-based Jaccard indices. To evaluate the differences in species composition and variance across sites between the two HTS, we performed a permutational multivariate analysis of variance (PERMANOVA) and the permutational analyses of multivariate dispersions (PERMDISP) with 10000 permutations, respectively. For the PERMANOVA and PERMDISP, we used the “adonis”, and “betadisper” functions in the “vegan” package ver. 2.5-6.

### Comparison of fish species detectability between eDNA metabarcoding and conventional methods

We evaluated species detectability between the two HTS by comparing the fish species lists of the two HTS with lists from conventional methods. Five sampling sites were selected from Kyushu and Chugoku districts (R23–27 in Fig.4). The fish fauna data obtained by conventional methods were based on the results of a previous study^43^. The conventional surveys were conducted through hand-net sampling and visual observation by snorkeling (see a previous study^43^ for the detailed methods). The count data of each species were replaced with the incidence-based datasets (presence or absence) for comparing with the eDNA metabarcoding datasets. Fish sequence reads of each sampling site obtained by eDNA metabarcoding were also replaced with the incidence-based data.

To test the detectability of species observed by conventional methods, the fish species compositions in five rivers were compared between the eDNA metabarcoding (iSeq and MiSeq) and the conventional methods. To visualize the differences in the species composition between HTSs and conventional methods, heat maps were illustrated for each sampling site. To assess differences in the number of species among methods at each river, the repeated measures analysis of variance (ANOVA) was performed among iSeq, MiSeq, and conventional methods. If a significant difference was found in repeated measures ANOVA, the Tukey-Kramer multiple comparison test was performed to analyze differences among methods.

Using fish faunal data obtained from iSeq, MiSeq, and conventional methods, the NMDS was performed in 1000 separate runs with Jaccard indices. The PERMANOVA was performed with 1000 permutations to assess the differences in fish fauna among the methods and sites.Furthermore, to evaluate variance across sites among methods, the PERMDISP was also performed with 1000 permutations. To visualize the number of species in each method and the number of common species between methods, Venn diagrams were illustrated for each river using the “VennDiagram” package ver. 1.6.2 in R^44^.

## Data availability

All data of the iSeq and MiSeq sequencing was shared in DRA (Accession number: DRA10593), and all used data, including all detected species by iSeq, MiSeq, and conventional methods, were shared in Supplementary Tables.

## Acknowledgments

We thank T. Kono and K. Yamaguchi for assisting in the field sampling. We also thank H. Fujii for assistance in the laboratory work and S. Miyazono for providing constructive comments on an earlier version of the manuscript. We thank K. Naka for assisting in iSeq running, and T. Kobayashi for running the MiSeq platform at the Illumina laboratory and for manuscript corrections. We also thank Illumina K. K. (Japan) for providing us with a sequencing cartridge kit for the iSeq run.

## Author Contributions

Y.A. and H.D. designed the study concept. R.I. and M.G. collected field samples, and M.G., H.D. and S.M. conducted a library preparation and iSeq sequencing. R.N., Y.A., S.M. and H.D. designed the methodology of data analyses and analyzed the data. R.N. and Y.A. led the writing of the manuscript, with critical contributions from all co-authors. R.N. prepared Fig. 1–4. All authors approved the final version of the manuscript.

## Additional Information

**Supplementary information** is available for this paper at Supplementary Figure S1 toS7, and Supplementary Table S1 to S10.

**Correspondence** and requests for materials should be addressed to R.N.

## References

1. Takahara, T., Minamoto, T., Yamanaka, H., Doi, H., & Kawabata Z. Estimation of fish biomass using Environmental DNA. PLoS ONE, 7, e35868 (2012).

2. Doi, H. et al. Environmental DNA analysis for estimating the abundance and biomass of stream fish. Freshw. Biol., 62, 30–39 (2017).

3. Stoeckle, M. Y., Soboleva, L., & Charlop-Powers Z. Aquatic environmental DNA detects seasonal fish abundance and habitat preference in an urban estuary. PLoS ONE, 12, e0175186 (2017).

4. Wu, Q. et al. Habitat selection and migration of the common shrimp, Palaemon paucidens in Lake Biwa, Japan—An eDNA-based study. Environmental DNA, 1, 54–63 (2019).

5. Tabarlet, P., Coissac, E., Hajibabei, M., & Rieseberg, L. H. Environmental DNA. Mol. Ecol., 21, 1789–1793. https://doi.org/10.1111/j.1365-294X.2012.05542.x

6. Barnes, M.A. & Turner, C. R. (2016). The ecology of environmental DNA and implications for conservation genetics. Conserv. Genet., 17, 1–17 (2012).

7. Jerde, C. L., Mahon, A. R., Chadderton, W. L., & Lodge, D. M. “Sight unseen” detection of rare aquatic species using environmental DNA. Conserv. Lett., 4, 150–157 (2011).

8. Thomsen, P. F. et al. Detection of a diverse marine fish fauna using Environmental DNA from seawater samples. PLoS ONE, 7, e41732 (2012).

9. Miya, M. et al. MiFish, a set of universal PCR primers for metabarcoding environmental DNA from fishes: detection of more than 230 subtropical marine species. Royal Soc. Open Sci., 2, 150088 (2015).

10. Yamamoto, S. et al. Environmental DNA metabarcoding reveals local fish communities in a species-rich coastal sea. Sci. Rep., 7, 40368 (2017).

11. Deiner, K. et al. Environmental DNA metabarcoding: Transforming how we survey animal and plant communities. Mol. Ecol., 26, 5872–5895 (2017).

12. Port, J. A. et al. Assessing vertebrate biodiversity in a kelp forest ecosystem using environmental DNA. Mol. Ecol., 25, 527–541 (2016).

13. Closek, C. J. et al. Marine vertebrate biodiversity and distribution within the Central California Current using environmental DNA (eDNA) Metabarcoding and Ecosystem Surveys. Front. Marine Sci., 16 (2019).

14. Ushio, M. et al. Environmental DNA enables detection of terrestrial mammals from forest pond water. Mol. Ecol. Resour., 17, e63–e75 (2017).

15. Ushio, M. et al. Demonstration of the potential of environmental DNA as a tool for the detection of avian species. Sci. Rep., 8, 4493 (2018).

16. Komai, T., Goto, O. R., Sado, T., & Miya M. Development a new set of PCR primers for eDNA metabarcoding decapod crustaceans, Metabarcoding Metagenom., 3, 1–19 (2019).

17. Thomsen, P. F., & Sigsgaard, E. E. Environmental DNA metabarcoding of wild flowers reveals diverse communities of terrestrial arthropods. Ecol. Evol., 9, 1665–1679 (2019).

18. Mychek-Londer, J. G., Balasingham, K. D., & Heath, D. D. Using environmental DNA metabarcoding to map invasive and native invertebrates in two Great Lakes tributaries. Environmental DNA. (2019).

19. Olds, B. P. et al. Estimating species richness using environmental DNA. Ecol. Evol., 6, 4214–4226 (2016).

20. Shaw, J.L.A. et al. Comparison of environmental DNA metabarcoding and conventional fish survey methods in a river system. Biol. Conserv., 197, 131–138 (2016).

21. Hänfling, B. et al. Environmental DNA metabarcoding of lake fish communities reflects long-term data from established survey methods. Mol. Ecol., 25, 3101–3119 (2016).

22. Deiner, K., Fronhofer, E. A., Mächler, E., Walser, J. C. & Altermatt, F. Environmental DNA reveals that rivers are conveyer belts of biodiversity information. Nat. Commun., 7, 12544 (2016).

23. Illumina. iSeq 100 sequencing system guide. Illumina https://support.illumina.com/downloads/iseq-100-system-guide-1000000036024.html (2019a).

24. Imamura, A., Hayami, K., Sakata, M.K., & Minamoto, T. Environmental DNA revealed the fish community of Hokkaido Island, Japan, after invasion by rainbow trout. Biodiversity Data Journal, 8, e56876 (2020).

25. Peters, L., et al. Environmental DNA: A New Low-Cost Monitoring Tool for Pathogens in Salmonid Aquaculture. Front. Microbiol., 9, 3009 (2018).

26. Tsukamoto, Y., Yonezawa, S., Katayama, N., & Isagi. Y. Detection of Endangered Aquatic Plants in Rapid Streams Using Environmental DNA. Front. Ecol. Evol., 8, 622291 (2021).

27. Illumina. Calculating percent passing filter for patterned and nonpatterned flow cells-A comparison of methods for calculating percent passing filter on Illumina flow cells. Illumina https://www.illumina.com/content/dam/illumina-marketing/documents/products/technotes/hiseq-x-percent-pf-technical-note-770-2014-043.pdf (2017).

28. Tsuji, S., Takahara, T., Doi, H., Shibata, N., & Yamanaka, H. The detection of aquatic macroorganisms using environmental DNA analysis—A review of methods for collection, extraction, and detection. Environmental DNA, 1, 99–108 (2019).

29. Fremier, A.K., Strickler, K.M., Parzych, J., Powers, S., & Goldberg, C.S. Stream Transport and Retention of Environmental DNA Pulse Releases in Relation to Hydrogeomorphic Scaling Factors. Environ. Sci. Technol., 53 (12), 6640–6649 (2019).

30. Jane, S.F., et al. Distance, flow and PCR inhibition: eDNA dynamics in two headwater streams. Mol. Ecol. Resour., 15(1), 216–227 (2015).

31. Colman, R.E. et al. Whole-genome and targeted sequencing of drug-resistant Mycobacterium tuberculosis on the iSeq100 and MiSeq: A performance, ease-of-use, and cost evaluation. PLoS Med. 16(4), e1002794 (2019).

32. Dohál, M., et al. Whole-genome sequencing and Mycobacterium tuberculosis: Challenges in sample preparation and sequencing data analysis. Tuberculosis, 123,101946 (2020).

33. Yamanaka, H. et al. A simple method for preserving environmental DNA in water samples at ambient temperature by addition of cationic surfactant. Limnol., 18, 233–241 (2017).

34. Doi, H. et al. The effects of ecosystem characteristics and species traits on species detection by eDNA metabarcoding compared to species records in Japanese lake fish communities. BioRxiv, preprint at https://doi.org/10.1101/2020.09.25.314336 (2020).

35. Uchii, K., Doi, H., & Minamoto, T. A novel environmental DNA approach to quantify the cryptic invasion of non-native genotypes. Mol. Ecol. Resour., 16, 415–422 (2016).

36. Edgar, R. C. Search and clustering orders of magnitude faster than BLAST. Bioinformat., 26, 2460–2461 (2010).

37. Takeuchi, A. et al. New PCR primers for metabarcoding environmental DNA from freshwater eels, genus Anguilla. Sci. Rep., 9, 7977 (2019).

38. Oka, S. et al. Environmental DNA metabarcoding for biodiversity monitoring of a highly diverse tropical fish community in a coral reef lagoon: Estimation of species richness and detection of habitat segregation. Environmental DNA, 3, 55–69 (2021).

39. Edgar, R. C. and Flyvbjerg, H. Error filtering, pair assembly and error correction for next-generation sequencing reads. Bioinformat., 31, 3476–3482 (2015).

40. Illumina. Cluster optimization overview guide. Illumina https://support.illumina.com/downloads/cluster-optimization-overview-guide-1000000071511.html (2019).

41. Ewing, B., Hillier L., Wendl M. C., & Green P. Base-calling of automated sequencer traces using Phred. I. Accuracy assessment. Genom. Res., 8, 175–185 (1998).

42. R Core Team. R: A language and environment for statistical computing. R Foundation for Statistical Computing, Vienna, Austria. https://www.R-project.org/ (2019).

43. Doi, H. et al. Evaluation of biodiversity metrics through environmental DNA metabarcoding compared with visual and capture surveys in river fish community. BioRxiv, preprint at https://doi.org/10.1101/617670 (2020).

44. Chen, H., & Boutros, P.C. Venn Diagram: a package for the generation of highly-customizable Venn & Euler diagrams in R. BMC Bioinformat, 12, 35 (2011).

45. Wickham H. ggplot2: Elegant Graphics for Data Analysis. Springer-Verlag New York. ISBN 978-3-319-24277-4, https://ggplot2.tidyverse.org (2016).

