## Supplementary Figures for "Illumina iSeq 100 and MiSeq exhibit similar performance in freshwater fish environmental DNA metabarcoding"

Supplementary Figure S1:

**Comparison of sequence quality between iSeq and MiSeq.**

Supplementary Figure S2:

**Relationship of sequence read per sample between iSeq and MiSeq after the merge pair step.**

Supplementary Figure S3:

**Relationship of sequence read per sample between iSeq and MiSeq after the quality filtering step.**

Supplementary Figure S4:

**Figure S4. Relationship of sequence read per sample between iSeq and MiSeq after the denoising step.**

Supplementary Figure S5:

**Figure S5. Relationship of remained sequence read per sample between iSeq and MiSeq for the taxonomic assignment.**

Supplementary Figure S6:

**Species accumulation curves of each samples in iSeq platform.**

Supplementary Figure S7:

**Species accumulation curves of each samples in MiSeq platform.**

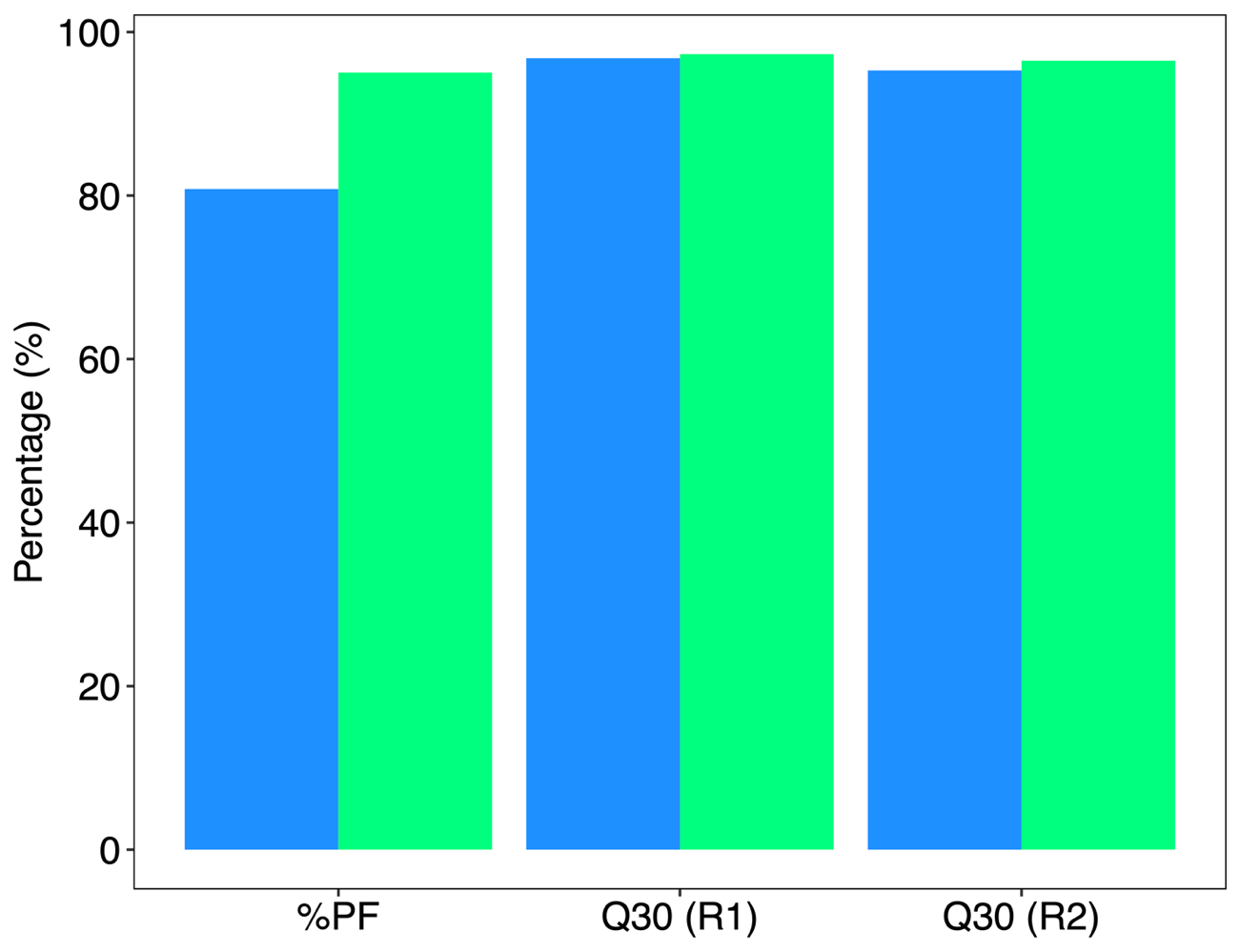

**Figure S1. Comparison of sequence quality between iSeq and MiSeq.** Blue and green bar plots show the results of iSeq and Miseq. Each bar also shows the percentage of pass filtering (80.8 vs. 95.1), Read 1 Q30 (96.8 vs. 97.3), and Read 2 Q30 (95.3 vs. 96.5), respectively. The bar plots were illustrated using “ggplot” function in ggplot2 package in R ver. 3.6.2.

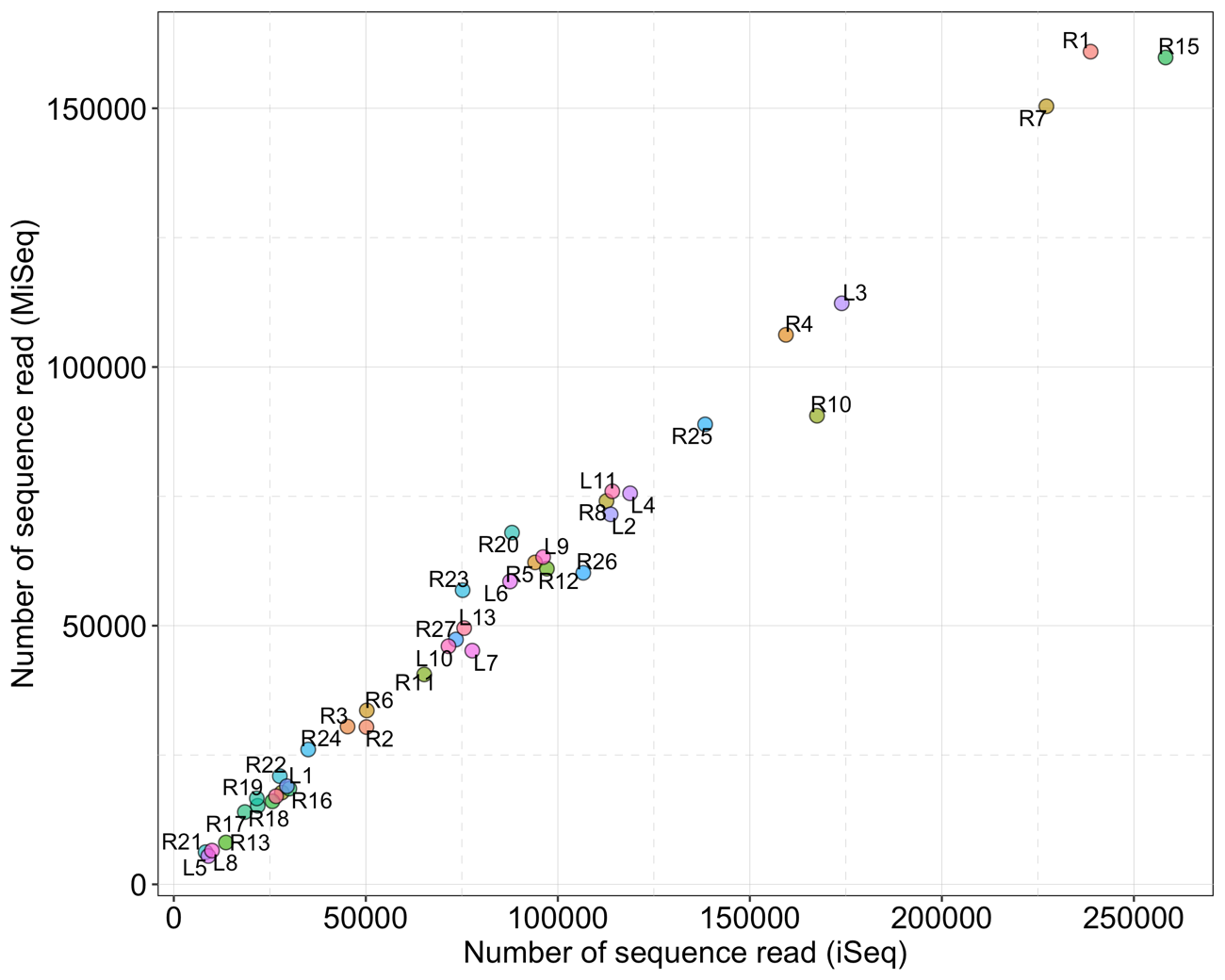

**Figure S2. Relationship of sequence read per sample between iSeq and MiSeq after the merge pair step.** The dot plot was illustrated using “ggplot” function in ggplot2 package in R ver. 3.6.2. There was significant positive correlation between the remained sequence reads of iSeq and MiSeq (spearman’s rank correlation, ρ = 0.991, *p* < 0.01)

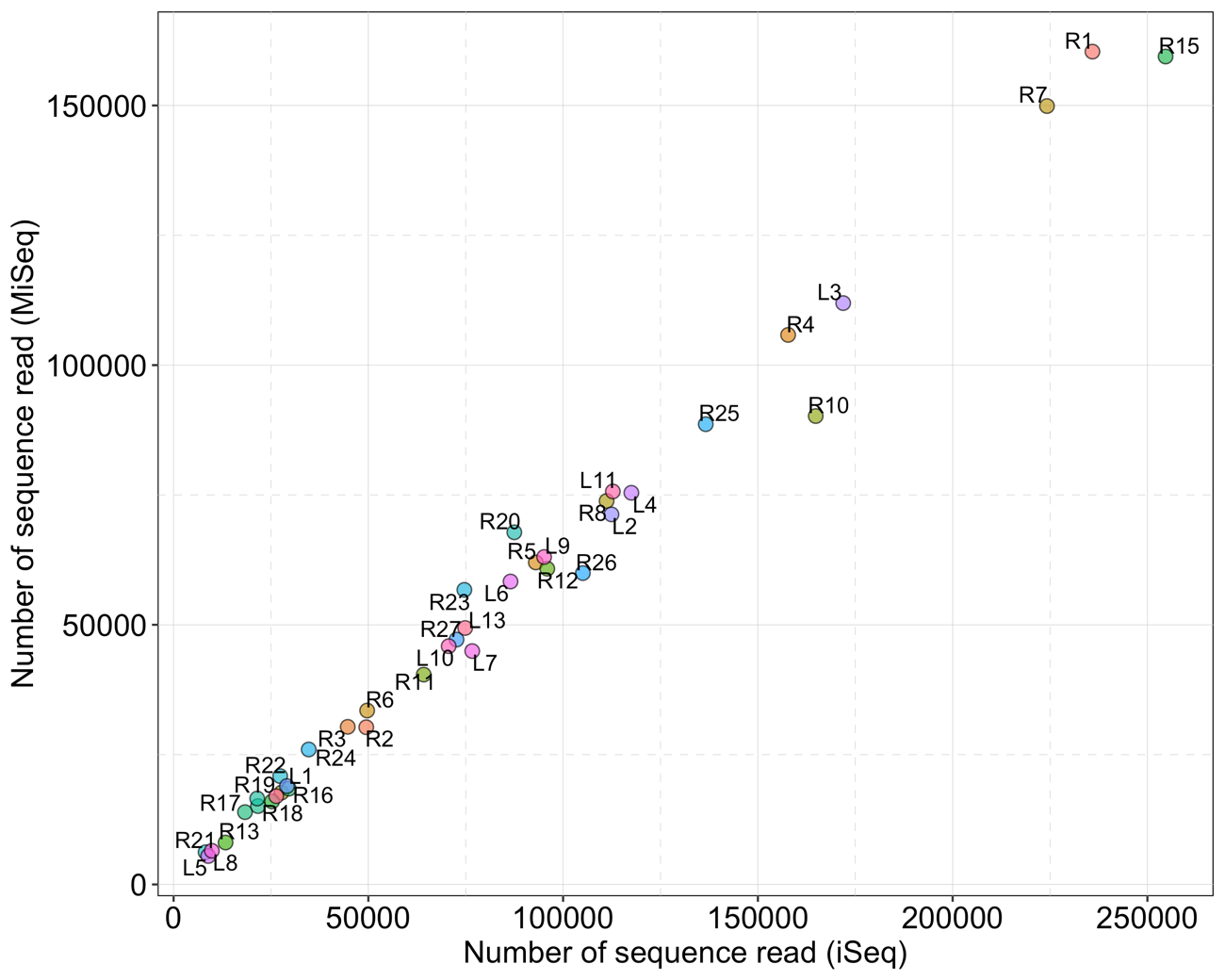

**Figure S3. Relationship of sequence read per sample between iSeq and MiSeq after the quality filtering step.** The dot plot was illustrated using “ggplot” function in ggplot2 package in R ver. 3.6.2. There was significant positive correlation between iSeq and MiSeq (spearman’s rank correlation, ρ = 0.991, *p* < 0.01)

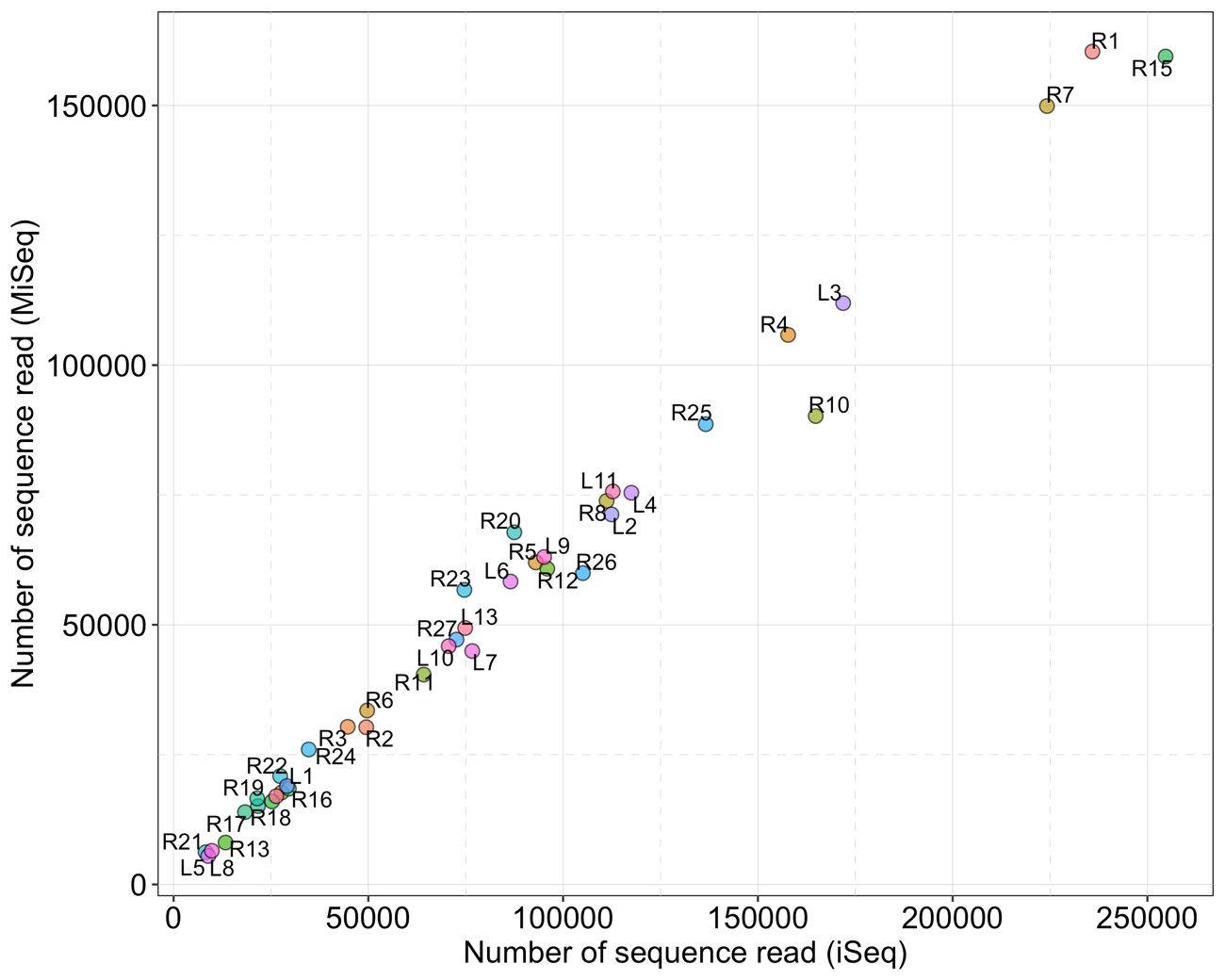

**Figure S4. Relationship of sequence read per sample between iSeq and MiSeq after the denoising step.** The dot plot was illustrated using “ggplot” function in ggplot2 package in R ver. 3.6.2. There was significant positive correlation between iSeq and MiSeq (spearman’s rank correlation, ρ = 0.993, *p* < 0.01)

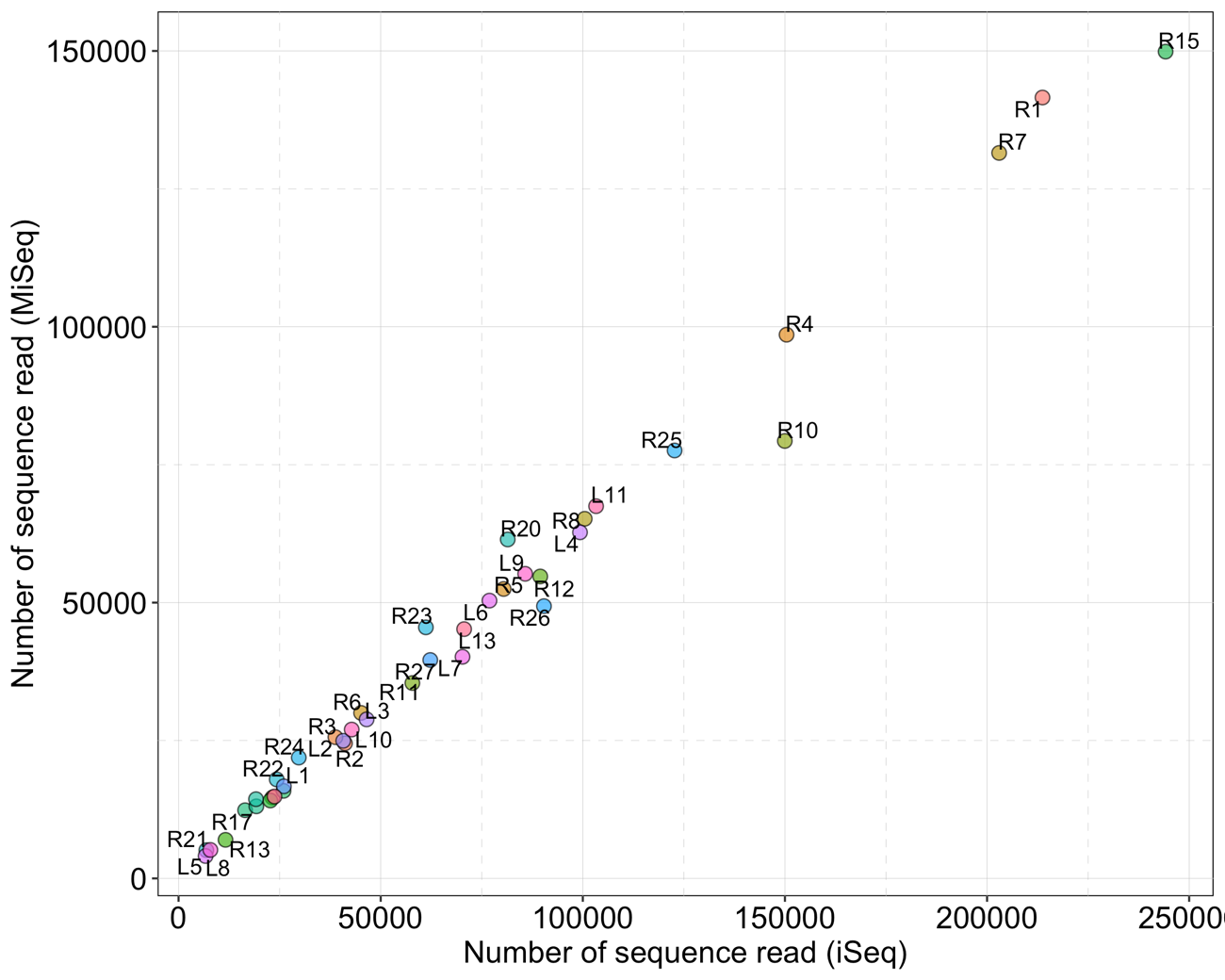

**Figure S5. Relationship of remained sequence read per sample between iSeq and MiSeq for the taxonomic assignment.** The dot plot was illustrated using “ggplot” function in ggplot2 package in R ver. 3.6.2. There was significant positive correlation between iSeq and MiSeq (spearman’s rank correlation, ρ = 0.993, *p* < 0.01).

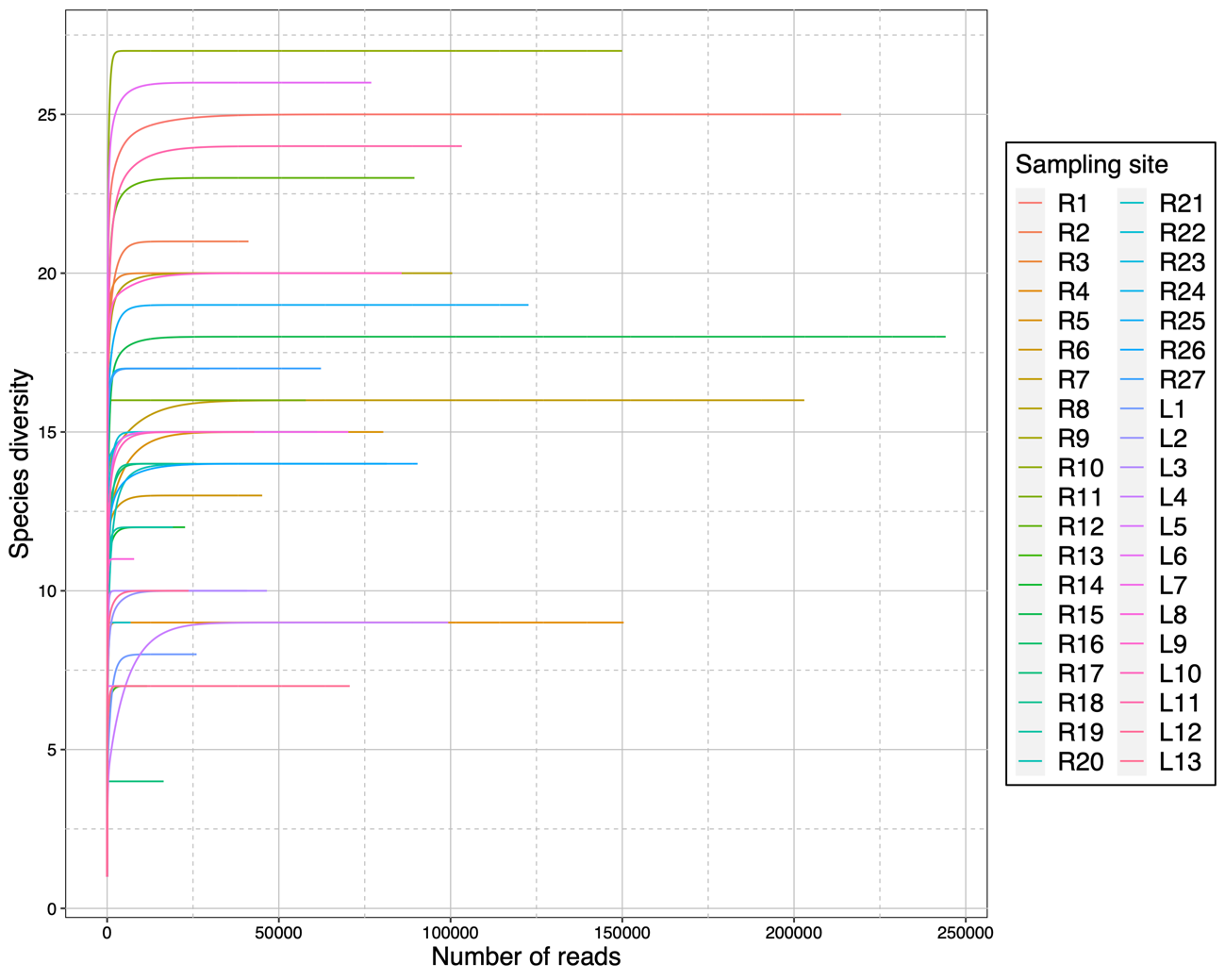

**Figure S6. Species accumulation curves of each samples in iSeq platform.** This graph was illustrated using “rarecurve” and “ggplot” function in vegan and ggplot2 package in R ver. 3.6.2, respectively.

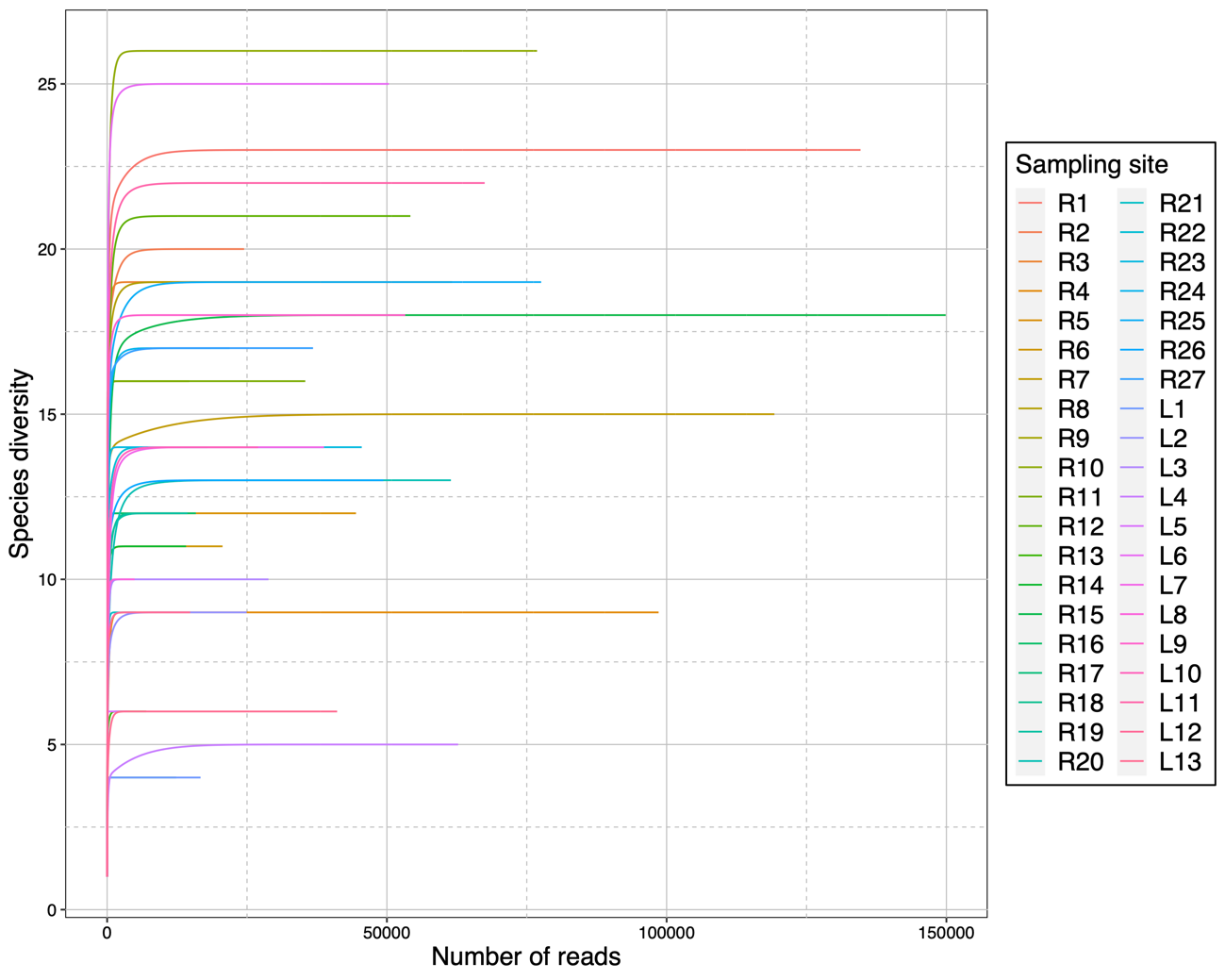

**Figure S7. Species accumulation curves of each samples in MiSeq platform.** This graph was illustrated using “rarecurve” and “ggplot” function in vegan and ggplot2 package in R ver. 3.6.2, respectively.
